# A dynamic control of human telomerase holoenzyme

**DOI:** 10.1101/525733

**Authors:** Mohammed E. Sayed, Ao Cheng, Gaya Yadav, Andrew T. Ludlow, Jerry W. Shay, Woodring E. Wright, Qiu-Xing Jiang

## Abstract

Human telomerase functions in maintaining genome stability by adding telomeric repeats to the termini of linear chromosomes. Past studies have revealed profound insights into telomerase functions. However, low abundance of functional telomerase and difficulty in quantifying its activity leave partially characterized its thermodynamic and kinetic properties. Using a newly developed method to count individual extension products, we demonstrate that human telomerase holoenzymes contain fast- and slow-acting catalytic sites. Surprisingly, both active sites become inactive after two consecutive rounds of catalysis. The fast active sites turn off ~40-fold quicker than the slow ones and exhibit higher affinity to substrates. In dimeric enzymes, the two sites work in tandem with the faster site functioning before the slower one. In monomeric enzymes, the active sites also perform single-run catalysis. Interestingly, the inactive enzymes can be reactivated by intracellular telomerase-activating factors (iTAFs) available in multiple cell types. Together, the single-run catalysis and the iTAF-triggered reactivation serve as a novel control circuit to ensure that the telomerase holoenzymes are dynamically controlled to match their number of active sites with the number of telomeres they extend. Such exquisite kinetic control of telomerase activity is expected to play important roles in cell division and ageing.

## INTRODUCTION

Telomeres refer to the terminal sequences of linear chromosomes in eukaryotic cells ^1^. Their catalytic extension provides an evolutionarily conserved solution to the “end replication” problem. To maintain proper telomeric length, eukaryotic cells utilize telomerase to catalyze addition of telomeric repeats using an intrinsic RNA template. In human cells, telomerase adds hexameric repeats, (TTAGGG)_n_, to chromosomal termini ^2^. During a cell cycle, a telomerase enzyme is recruited to a transiently-uncapped telomere before it can function in a controlled fashion ^3^. It is believed that the telomerase preferentially acts on shorter telomeres ^4–8^. Every telomere should be acted on in order to maintain proper telomere length equilibrium; otherwise, some telomeres would become shorter over time, leading to cellular senescence ^9^. How telomerase-expressing cells regulate telomere length in a global scale remains unclear.

Telomerase plays a critical role in human diseases, in particular cancers and other age-related diseases. With down-regulated telomerase activity, differentiating cells reach a critical state after a certain number of cell divisions such that the cell cycle will be arrested and cells will enter “senescence”. How a cell senses changes in telomere length before senescence remains a mystery. Approximately 85-90% of human cancers exhibit elevated telomerase activity ^1, 10–18^. The importance of telomerase activity to cancer cells and other proliferative stem-like cells has been well demonstrated. Chemical inhibitors and activators of human telomerase are being explored for cancer treatment and anti-ageing therapies, respectively ^19–21^. These two aspects make the structural and functional studies of human telomerase and telomere maintenance a key area in cell biology and cancer biology ^22–30^.

In human cells, the telomerase holoenzymes are heterogeneous and contain both monomers and dimers ^31^. A dimeric holoenzyme contains two copies of hTERT (**t**elomeric **r**everse **t**ranscriptase), dyskerin and hTR (**T**elomerase **R**NA **c**omponent) as well as other factors. It has an apparent mass of ~700 kDa ^12, 32, 33^. A monomeric telomerase holoenzyme has one copy of hTERT, hTR, and TCAB1, and two copies of NHP2, dyskerin, NOP10, and GAR1 as recently resolved by cryoEM ^34^, and is of similar size as the dimeric form. Each monomer has one active site. By reconstitution in cultured cells, every hTERT/hTR pair forms a minimally active enzyme that has one active site ^35^. A proliferating human cell may have 50-100 copies of functional enzymes ^9, 36^. It remains unclear how free telomerase complexes are physically recruited to recognize the telomeres when chromosomal 3’-overhangs become accessible (uncapped) during the S phase; Nor is it well understood how a telomerase adds telomeric repeats in a processive fashion or finishes the reaction when the enzyme stochastically falls off its product ^9, 37–41^. Energetically, telomerase uses the chemical energy from dNTP hydrolysis to catalyze its RNA-guided DNA synthesis. For every six-nucleotide repeat, perhaps following a Boltzmann distribution among the states with different numbers of nucleotides added, the DNA-RNA hybrids are presumably most stable when all six positions are paired, explaining the main peaks for every repeat addition observed by gel-based activity assays. For processive extension, an energetically costly step is expected to melt the DNA-RNA hybrid in order to translocate the substrate by six nucleotides or fall off from the product sporadically to terminate the reaction. The kinetics and the mechanistic programs for the processive addition of telomeric repeats by telomerase are yet to be elucidated.

Prior studies of the human telomerase holoenzyme have revealed important insights ^12, 42, 43^, but the dynamic control for its catalytic activity, the structural relationship between hTR and hTERT subunits and the interactions between the telomerase holoenzyme and its substrates remain incompletely understood. Semi-quantitative analyses by gel-based direct or TRAP assays have derived incomplete kinetic and thermodynamic properties of telomerase, partially because low abundance of telomerase has significantly limited quantitative analysis. In this study, we will use quantitative analysis to reveal an unexpected kinetic property of human telomerase.

## RESULTS

### In single-turnover settings, human telomerase becomes inactive after two sequential runs of catalysis

In order to analyze telomerase activity quantitatively, we took advantage of telomerase’s ability to stably bind telomeric repeats (TTAGGG)_3_ (R3) at the GGG position, but quickly fall off when it adds three nucleotides to the substrate and reaches the TTA position *in vitro* (**Fig. 1A**) ^32^. *Each addition of TTA is thus half a telomeric repeat and constitutes a single-turnover condition for analyzing the catalytic activity*. A single-step pull-down using triple telomeric repeats as ligands enriched active enzymes directly from cell lysates (Supplementary **Fig. S1A**), which contained both monomers and dimers ^31^. When studying the macroscopic, thermodynamic behavior of telomerase, we were not able to computationally separate the monomers from the dimers as what was done during cryoEM analysis or as in single molecule enzymology ^34, 44^. Instead, we used two sequential pull-downs to separate the dimers from the monomers. A conventional TRAP assay, using lysates from 50, 500 and 5000 cells as a control, showed that a single-step pulldown sequestered approximately 25% of the total activity (Supplementary **Fig. S1A**), which is not surprising because only enzyme molecules with their active sites accessible to substrates were pulled down.

**Figure 1:**
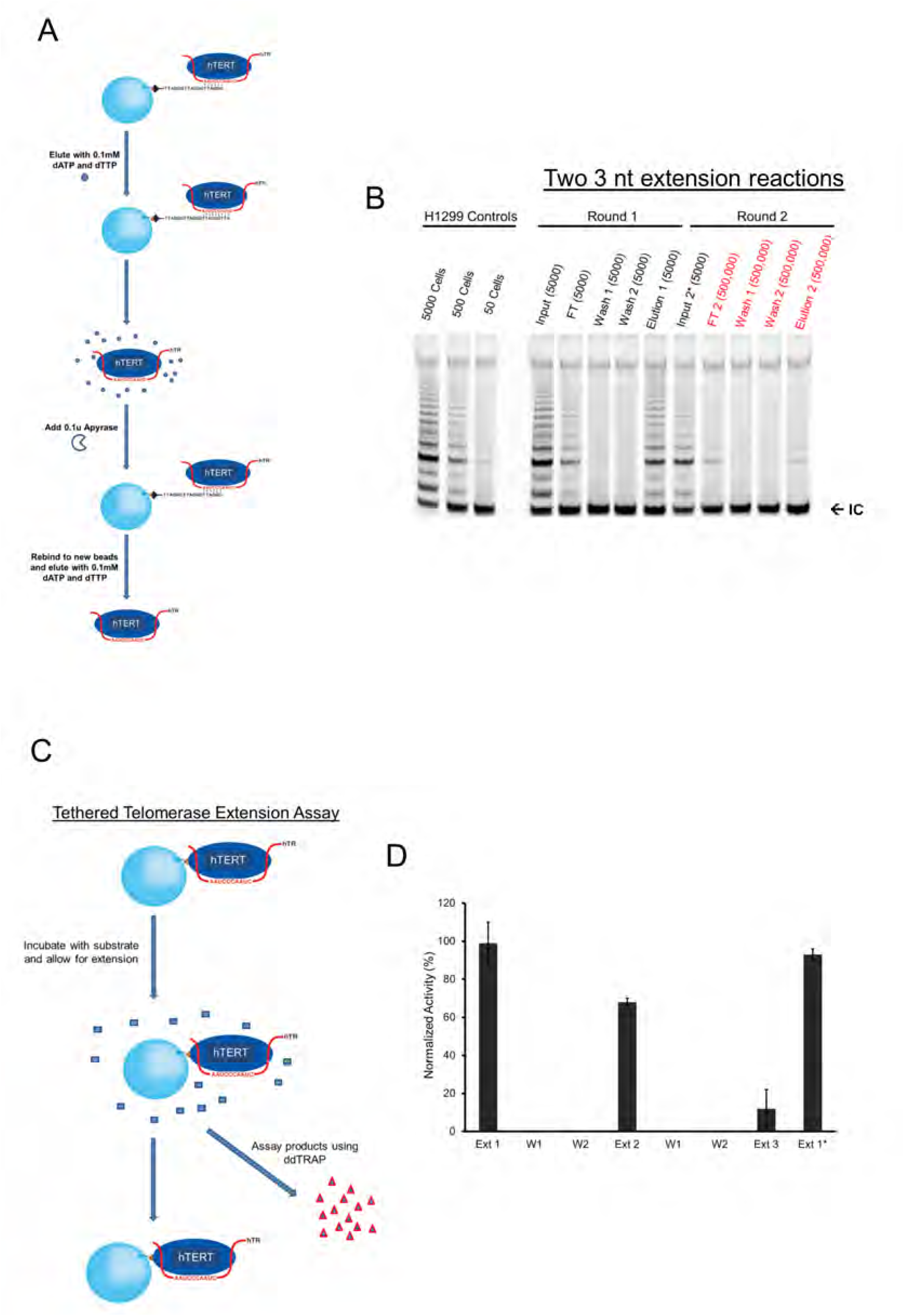
Catalysis dependent loss of telomerase activity. ***(A)*** Schematic representation of two repetitive cycles of affinity-based pull-down and elution from R3 telomeric substrates in order to purify endogenous telomerase from H1299 cells. Fresh beads and oligos were used for the second round. dATP and dTTP were inactivated by apyrase enzyme in order for eluted telomerase to bind to fresh beads. ***(B)*** Gel-based TRAP assay of samples from the telomerase going through double affinity pull-down. Samples were loaded according to cell equivalence. Cell equivalence is labeled separately (50, 500, 5000 etc.). The bottom bands (IC) represent the internal PCR control, iTAS. H1299 control samples on far left represent total activity from cell lysate. ***(C)*** Schematic representation of the extension assay of tethered telomerase through the biotinylated hTERT. After every reaction cycle, extension products were separated from the beads (and the enzymes) and quantified using the ddTRAP assay. ***(D)*** ddTRAP assays of the tethered telomerase. Ext 1 is the first reaction. The enzymes were washed twice before the second reaction (Ext 2). Two more washes before the third reaction (Ext 3). “Ext 1*” denotes the control sample with the delayed extension reaction (after being kept for 4 hours at room temperature before substrates were presented to it). Error bars: *s.d*. (n=3).

Our data also show that the gel-based TRAP assay is insensitive to changes of telomerase activity by less than five folds. The direct assays based on radioactivity suffered from similar insensitivity and non-linearity. To increase the amount of telomerase for experiments, we generated a stable cell line overexpressing both hTR and N-terminally biotinylated hTERT (Supplementary **Fig. S2A**). When compared with parental H1299 cells and stable H1299 cells overexpressing hTERT alone by the gel-based TRAP assay, the engineered cells had ~20 fold more activity (Supplementary **Fig. S3A**), and will be called the *super H1299*. The biotinylation was accomplished by the mammalian biotinylation machinery inside the cells ^45^. When incubated with streptavidin-coated beads, the biotinylated telomerase bound to the beads, but the endogenous enzyme did not (supplementary **Fig. S2B** vs. **S2C**), suggesting that the recombinant hTERT, ~10 kDa heavier than the endogenous one in SDS-PAGE, was indeed biotinylated. Biotinylation provided an effective way to selectively enrich recombinant telomerase, and allowed the separation of the recombinant hTERT from the endogenous hTERT as well as possible contaminating proteins (supplementary **Fig. S3B-C**).

We used a two-step pull-down procedure in order to enrich selectively endogenous telomerase holoenzymes (**Fig. 1A**). Between the two steps, apyrase was introduced to remove dATP/dTTP from the eluate of the first pulldown (Supplementary **Fig. S4**). An aliquot of the sample loaded to the substrate (R3) beads, equivalent to the enzymes from 5,000 parental H1299 cells, was analyzed (Input in **Fig. 1B**). Equivalent samples (cell equivalency) from other fractions were analyzed as well. A small amount of activity (<10%) showed in the flow-through (FT in **Fig. 1B**). After two washing steps, the eluted fractions showed strong activity (~25%; Elution 1 in **Fig. 1B**). Based on the test results in supplementary **Fig. S4**, 0.1 units of apyrase were introduced to hydrolyze dTTP and dATP in the Elution 1 before the eluate was mixed to fresh R3 beads (Input 2 in **Fig. 1B**). Surprisingly, although we expected to recover ~25% of the loaded telomerase activity in the second eluate, literally no activity was detected in all subsequent fractions (FT2, Wash 1 & 2, and Elution 2 in **Fig. 1B**), suggesting that after binding to fresh substrates and / or performing the second round of 3-nucleotide extension, telomerase became inactive (**Figs. 1A & 1B**). To highlight this point, 100 x more materials, equivalent to those from 500,000 cells, for all fractions from the second pull-down were analyzed in the gel-based TRAP assay [Elution 2 (500,000) in **Fig. 1B**].

The surprising results in **Fig. 1B** suggested the possibility that the telomerase holoenzyme, once separated from the other components of the cell lysates, shuts itself off after *two runs of short 3-nucleotide single-turnover extension reactions*. To examine if this observation stemmed from the apyrase treatment or the short single-turnover extensions, we tethered biotinylated telomerase from super H1299 cells to streptavidin-coated beads and used a more quantitative digital droplet TRAP assay (ddTRAP) to count individual extension products. *ddTRAP is accurate to one extension product, and is much more sensitive than both the direct methods and the conventional TRAP assays. It is linear in a broad dynamic range of the extension products being detected in our experiments* ^46^. On the beads, streptavidin molecules were at least 50 nm apart such that the telomerase holoenzymes tethered on the beads could not physically interact with each other. The ddTRAP assay counted individual extension products in an all-or-none fashion, only registering the frequency of successful catalytic interactions between the enzymes and the DNA substrates (TS primers) without measuring the length of the products. Careful comparison has found that within the proper concentration ranges of the reactants, the results of ddTRAP with those are linearly proportionate with the conventional TRAP assay that counts the length of processive reactions^47^. The ddTRAP is more precise in quantification and fast for high-throughput analysis, and thus suitable for our analysis.

When analyzed in a time-lapsed fashion, the tethered telomerase was stable for at least 4-5 hours at room temperature (**Fig. 1C and Fig. 2A-B**), suitable for the kinetic analysis. After each run of processive extension, longer than the half-turn addition of 3-nucleotides in **Fig. 1A**, the products were separated from the enzymes and quantified (**Fig. 1D**). During the second run (Ext 2 in **Fig. 1D**), the reaction products were 60-70% of those from the first run (Ext 1). During the third run (Ext 3), the yield dropped to < 10% of those in the first run with the same amount of tethered enzyme. As a control, the same enzyme preparation going through the same mechanical manipulations for three rounds without seeing the telomeric TS primers (DNA substrates) in the first two runs generated nearly the same amount of products (Ext 1* in **Fig. 1D**) as the first run (Ext). This indicated that the waiting time and the manipulations through tethering and washing caused no substantial loss of activity in our assays. The loss of activity is thus *use-dependent*.

**Figure 2:**
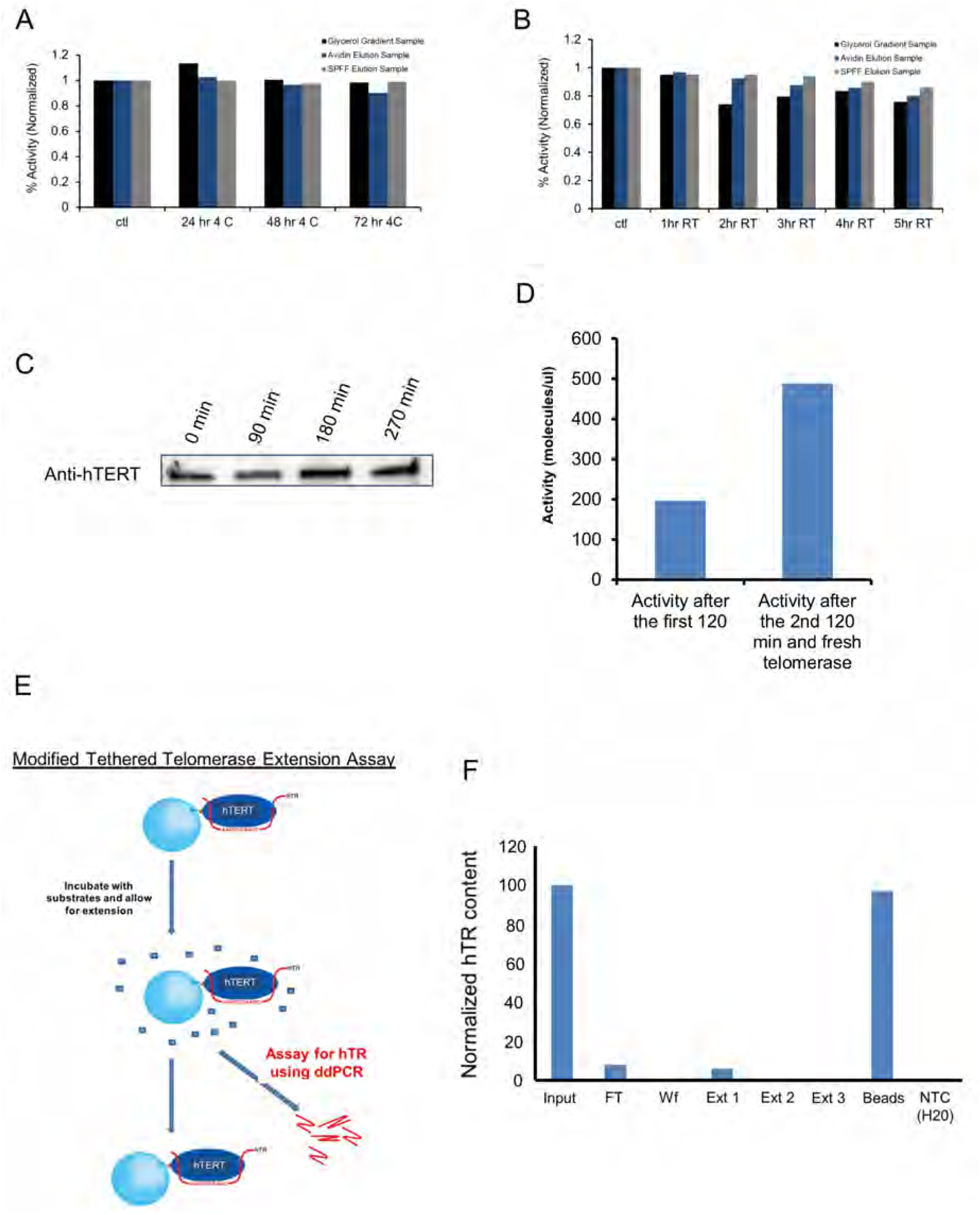
Activity loss of telomerase was not due to enzyme instability, protein degradation, product inhibition or loss of hTR from complex. ***(A)*** Partially purified telomerase (at different stages indicated by colored bars) retained most activity over 3 days at 4° C as quantified by ddTRAP activity assay. ***(B)*** Partially purified telomerase enzyme over 5 hrs at room temperature, RT, as quantified by ddTRAP activity assay. ***(C)*** Western blot of hTERT showing telomerase stability throughout the full cycle of the extension assay of the tethered telomerase (nearly 5 hours at RT). ***(D)*** No product inhibition of telomerase activity. After one extension reaction for 120 minutes, the enzyme was separated from the reaction mixture containing the extension products. Fresh telomerase (equal amount) was added to the same reaction mixture. After another 120 min of extension, the products (from two tandem reactions) were quantified with ddTRAP activity assay (right), roughly twice of the control (left) from the first reaction. ***(E)*** Schematic representation of modified tethered telomerase extension assay. In this assay, ddPCR was used to quantify the hTR content in the products to determine whether hTR dissociated from the telomerase complex. ***(F)*** ddPCR quantification of hTR during multiple rounds of extension reactions (Ext 1 - 3). >95% of hTRs remained bound to the tethered enzymes.

The loss of telomerase activity after two single-turnover runs of catalysis might result from multiple possibilities. To name a few, telomerase may be degraded or lost during the apyrase treatment, its RNA component (hTR) might dissociate from the complex, one or more of its intrinsic accessory factors might fall off after catalysis, or the products from its extension reaction might cause inhibition even in the presence of a saturating concentration of substrates. We tested all of these possibilities. First, partially purified telomerase using streptavidin-beads or other methods were stable for a few days at 4 °C (**Fig. 2A** and Supplementary **Figs S5A-C**) or for 5-6 hours at room temperature (**Fig. 2B**) with the loss of activity limited to 5-15%. The endogenous telomerase directly from cell lysates, which also shut off after two runs of extensions, lost a bit more activity (35%) over a period of 5 hours (Supplementary **Fig. 1B**), but it is much lower than the > 90% use-dependent loss. We thus preferentially used partially purified enzymes for our studies.

Second, western blot of both endogenous and recombinant hTERT found that the tethered enzymes after two rounds of extension reactions (**Fig. 1D**) suffered from no obvious hTERT degradation (**Fig. 2C**). Third, RT-qPCR analysis of the hTR content found that the “Elution 2” fraction in **Fig. 1B** had ~25% of the hTR in the Elution 1 as we expected, but ZERO activity in even 100 x more materials (**Fig 1B** & supplementary **Fig. S6**). Fourth, using the recombinant telomerase containing the N-terminally biotinylated hTERT (supplementary **Fig. S3A** and **Fig. 1C**), we started one reaction with a batch of fresh enzyme and after 2 hours, separated the enzymes from the products. When fresh telomerase was added into the reaction mixture that contained the products from the first round of catalysis, it produced a similar amount of products (the total product amount doubled as in **Fig. 2D**). No product inhibition existed for telomerase.

Fifth, when ddPCR was used to detect reverse-transcripts of hTR extracted from the tethered enzymes (**Fig. 2E**), almost all hTR was retained with the tethered enzymes, suggesting that the tethered telomerase holoenzyme did not fall apart. Last, when the holoenzymes after multiple manipulations were blotted for dyskerin, NHP2 and NOP10, all three accessory proteineous factors of the active enzymes were retained (**Fig. S5D**), suggesting that the key protein factors were still retained because the enzyme did not fall apart, although we could not rule out thatan unknown factor fell off the holoenzyme. These data revealed the integrity of a telomerase RNP complex by retaining the core components of hTERT, hTERC and key protein factors (**Fig. 2F**). Together these experiments demonstrated that the use-dependent loss of telomerase activity was not caused by chromatographic manipulations, apyrase treatment, instability or degradation of the holoenzyme, hTR dissociation, release of key accessory factors or product inhibition.

Uses-dependent loss of activity is not unusual. Conventional “single turnover” enzymes are good examples. With saturating substrates, a normal enzyme catalyzes a reaction continuously with a linear increase of products over a long period of time (red dashed lines in **Figs. 4A-B, 4E-F**). A single-turnover enzyme, however, is self-limiting by turning itself off after one round of catalysis. In a similar fashion, human telomerase shuts off after two runs of single-turnover processive extension. At the molecular level, there are three potential explanations. One is that each enzyme, e.g. monomeric telomerase, has only one active site, which has the same affinity to the substrates, and is able to count the rounds of its catalysis and shut itself off after exactly two rounds. As a thermodynamic system, such counting in an exact fashion is improbable. The second possibility is that there are two types of monomeric enzymes, such as monomeric telomerase holoenzymes made of the core factors and different factors and named as fM_1_ and sM_1_, whose active sites differ. The fM_1_ functions mainly in the first round of catalysis and the sM_1_, a good fraction of which does not bind to the TS primers in the first round, works in the second. Third, there are two types of active sites in a dimeric enzyme, both of which shut off after two separate reactions. The dimeric enzymes may co-exist with fM_1_ or both fM_1_ and sM_1_ especially in consideration of the mixture of dimers and monomers seen by cryoEM ^34^.

### Human telomerase holoenzyme from super H1299 cells is heterogeneous in composition

In literature, the telomerase holoenzymes in human cells are heterogeneous, containing both dimers and monomers ^32, 48–50^. Minimally active hTERT/hTR complex is monomeric with only one active site and simple kinetic behavior ^35^; but without other protein components of the holoenzyme (Supplementary **Fig. S5D**), it may be absent of real physiology. Enzymes attached to slides or exposed to or near air-water interfaces were found to be a mixture of both dimers and monomers. It is unclear whether such operations might change the ratio of dimers to monomers ^31^. The acutely assembled monomeric enzymes analyzed by cryoEM showed a complex of ~590 kDa. To evaluate our recombinant enzymes, we used a continuous glycerol gradient to separate active enzymes from cell lysates. Thyroglobulin (~669 kDa), a molecular weight marker, was used as a landmark in our gradient (**Fig 3A**, fractions F7/F8). Western blotting found that the fractionated enzymes from the super-H1299 cells were distributed near the bottom of the density gradient (F8-F11), suggesting that the holoenzymes are quite heterogeneous in size, apparently heavier than a 669 kDa globular protein (**Fig. 3B-C**). Here we did not take consideration of variations in partial specific volumes, frictional resistance, and solution viscosity and density for individual proteins because they are difficult to measure for each complex individually. Instead, we used a molecular weight marker. Similarly, size-exclusion chromatography of the glycerol-gradient fractions in a Superose 6 column found that a major fraction of the telomerase had a retention volume of 11-19 ml, equivalent to globular proteins in a broad range of ~0.3-0.9 MDa (**Fig. 3D**). The gel-based TRAP assay showed that the active human telomerase holoenzymes may be larger or smaller than the thyroglobulin (669 kDa) in size. Even though 700 kDa dimers and the 590 kDa monomers were not separated well in the density gradient or by size-exclusion chromatography, the active enzymes isolated from the super H1299 cells show heterogeneous composition containing both monomers and dimers as reported before ^34 31, 51^.

**Figure 3:**
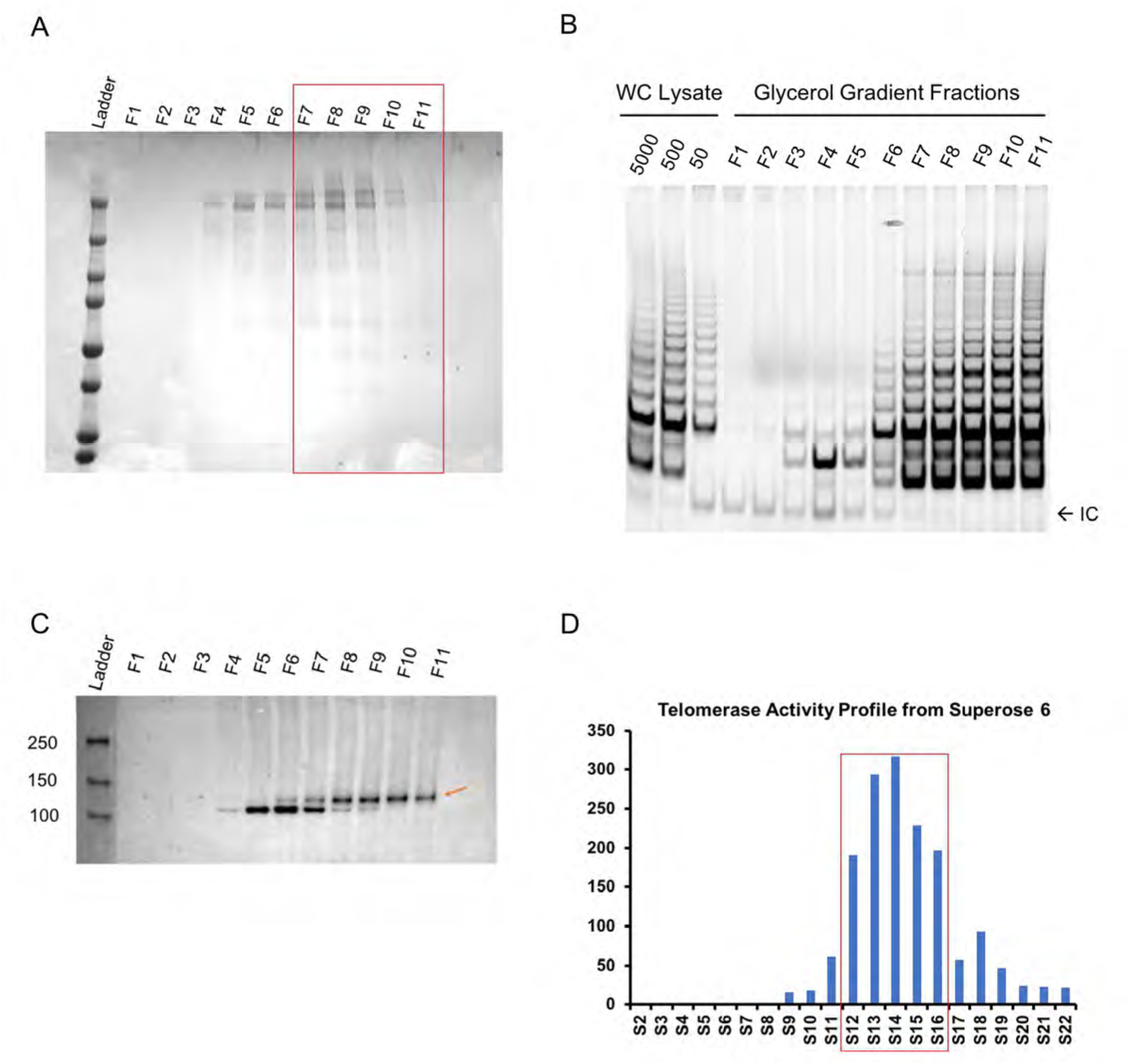
Recombinant telomerase complexes from super H1299 cells have a major fraction of dimers. Molecular weight marker thyroglobulin (~669 kDa) was run on a 10-30% glycerol gradient in parallel with recombinant telomerase. ***(A)*** Coomassie-stained SDS-PAGE assay of thyroglobulin in all eleven gradient fractions (top F1 to bottom F11). Red box marks the fractions where telomerase activity was detected. ***(B)*** Western blot (anti-hTERT) of gradient fractions shows that the recombinant hTERT (red arrow) is primarily in F8-F11. The holoenzyme is thus slightly heavier than 669 kDa. ***(C)*** Recombinant telomerase run through a Superpose 6 column. ddTRAP assay of the eluted fractions is plotted against fraction # (1 ml each). Red box indicated the major activity containing fractions. Thyroglobulin run on the same column eluted with its peak in fraction S14-S15, suggesting that the recombinant telomerase holoenzyme complex be heavier than 669 kDa. ***(D)*** Gel-based TRAP assay on gradient fractions. Lysates of super H1299 fractioned in the glycerol gradient were assayed. Majority of the activity was found in fractions 8-11.

### Time-lapse experiments reveal two distinct active sites for human telomerase

Partially fractionated telomerase holoenzymes were sufficiently stable for measurements in a period of 3 to 6 hours (**Fig. 2A**). The same was found to be true for telomerase enriched in glycerol-gradient fractions of cell lysates (**Fig. 3D**), the fractions eluted from single-step pulldown (**Fig 1B**), and the fractions of the biotinylated enzymes eluted from the monomeric avidin beads (**Fig. 3B**). High stability of the active telomerase in all these preps made it feasible to conduct the time-lapse experiments under different conditions.

We asked the question of whether the active sites in a dimeric telomerase holoenzyme or in monomeric enzymes were both active at the same time and if both active, whether the two sites would both undergo catalysis-dependent loss of activity. We first characterized the basic thermodynamic and kinetic properties of the active sites. Because two different monomers may be kinetics similar to dimers with two different active sites, our analysis will use dimeric enzymes as examples and the principles then apply to the two different monomers. A dimeric enzyme with two types of active sites might be in three distinct conformational states: the pristine enzyme (E_0_) with both active sites functional, the once-used enzyme (E_1_) with one site functional and the other nonfunctional, and the exhausted enzyme (E_2_) with no activity. A monomeric enzyme has one active site in either the active (M_1_) or the exhausted state (M_2_). Quantitative kinetic analysis would be needed in order to characterize enzymes distributed among these states ^47^.

With saturating concentrations of substrates (200 nM), catalytic activity of the telomerase from H1299 cell lysates or partially purified in a continuous glycerol gradient was compared with the enzymes eluted from (TTAGGG)_3_-conjugated beads (as the Elution 1 in **Fig. 1B**). The same amount (cell equivalency) of enzymes were used for extension reaction, but stopped at different time points by heat inactivation to halt the reactions and dissociate the products (**Figs 4A, 4B**). The released products were quantified by ddTRAP and normalized against the total products at 90 minutes. Without exception, the enzymes from different preparations all exhibited a fast and a slow kinetic component (**Figs 4A-4D**; supplementary **Fig. S7**). The fast one was saturated after ~5 minutes. The slow one slowly increased without saturation at 90 minutes, but with a longer time, it did saturate after ~300 minutes (**Figs 4A-B, 4E-F** & **S7**). The kinetic difference between the two components is striking (**Figs 4A, 4E**). Both components deviate significantly from what is expected from enzymes that are continuously active (red dash-lines based on the initial reaction rates in **Figs 4 & S7**). In accord with the use-dependent loss of activity, the two saturating kinetic components in product accumulation suggest that there are two different types of active sites that both become inactive after catalysis. As depicted above, the two kinds of active sites may come from two distinct monomeric holoenzymes or be contained within one dimeric complex.

**Figure 4:**
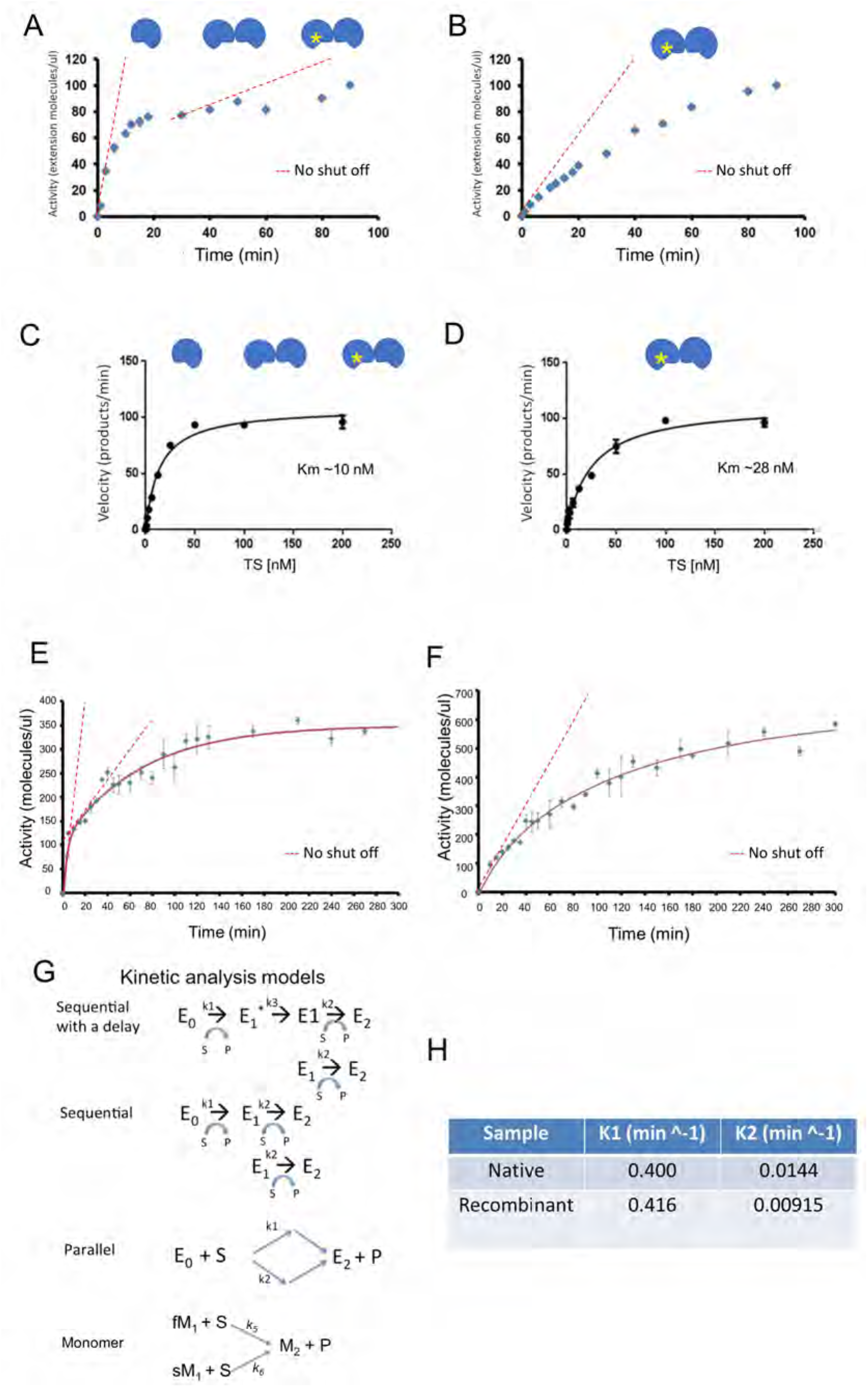
Two distinct active sites in human telomerase. ***(A)*** Time course for endogenous telomerase before one-step pull-down purification as in **Fig 1B**. Telomerase cartoons indicate three populations within the sample. Yellow star indicates a “once used” enzyme. The enzyme without a star represents either a “pristine” E0 dimer or an M1 monomer. The red dashed lines represent the amount of products if the active sites remained continuously active. ***(B)*** Time-course of endogenous telomerase after one-step pull-down purification (Elution 1). The telomerase cartoons indicated a “once-used” E1 dimer and slow M1 monomer, respectively. The red dashed line represents the amount of products if the active sites remained continuously active. ***(C)*** Michaelis-Menten plot for fast-acting components of the endogenous telomerase in panel **A**. Fitting of the data found *K_m_* ~10 nM. ***(D)*** Michaelis-Menten plot for endogenous telomerase used in panel **B**. *K_m_* ~ 28 nM. ***(E)*** Time course for ddTRAP assay of partially purified endogenous telomerase activity in fractions from glycerol gradient. Error bars: *s.d*. (n=3). The red dashed lines represent the amount of products if the active sites remained continuously active. ***(F)*** Time course for ddTRAP assay of partially purified recombinant telomerase in glycerol gradient fractions. Error bars: *s.d*. (n=3). The dashed line represents the amount of products if the active sites remained continuously active. ***(G)*** Parallel Markov models for monomeric telomerase by switching fast- (fM_1_) and slow-acting (sM_1_) monomers into inactive monomers (M2) and three different kinetic models for a telomerase dimer: Sequential with delay, Sequential and Parallel. ***(H)*** “Fast” and “slow” kinetic constants (*k*_1_ and *k*_2_) from data in panels E and F after two-exponential fitting.

Compared to the two kinetic components of the endogenous telomerase, the telomerase from the one-step single-turnover affinity purification using the R3-beads exhibited *only the slow component* (**Fig. 4B**). Because the single-turnover affinity purification would remove all monomeric enzymes (fast or slow), the leftover enzymes must be dimeric (**Fig 4B**), which has the same kinetic constant as the slow-component in the mixed enzymes from cell lysate (**Fig. 4B vs 4E**). Comparing this result with those in **Figs 1B** & **4A**, we deduced that after the first single-turnover addition of 3-nucleotides, the fast kinetic components (**Fig. 4A**) either dimers in E_0_ state or fast monomers fM_1_ and the slow components (monomers in sM_1_ and dimers in E_1_), which remained bound to the beads or were able to add 3-nt and elute out, were removed such that the Elution 1 fraction (**Fig. 4B**) was dominated by the dimeric enzymes with slow-acting active sites (E_1_). The two kinetic components were different in time domain such that two exponential components were needed to fit the data (**Figs 4E-F**). As compared in supplementary **Fig. S7**, one exponential component is not sufficient to fit the data.

We next evaluated the initial rates of product accumulation by varying the DNA-substrate concentration while keeping the dNTPs saturated, under which the binding of the DNA substrate became rate-limiting. The average initial rate, which was represented by the slope within the first five minutes and extrapolated to near t = 0 min (the boundary condition), was obtained for the fast component as illustrated in **Fig. 4A**. The average initial rate in the first 10 minutes was obtained for the slow-acting component as showed in **Fig. 4B**. We normalized the initial reaction rates against the maximal rates at 200 nM substrates and plotted them against substrate concentration (**Figs 4C & 4D**, respectively). Without calibrating the absolute quantity of the telomerase, we instead used the same amount of enzyme fractions for the reactions stopped at different time points and repeated the experiments in triplicates. Because we measured the initial rates with a trace amount of enzymes and a small fraction of product accumulation, a Michaels-Menten equation (*V_0_ / V_max_ = 1 / (1+[S] / K_m_)*) could be used to describe the normalized initial rate as a function of the DNA substrate, yielding *K_m_* = 10 nM and 28 nM for the fast-acting and slow-acting components, respectively. These are fairly close to what were measured before using different DNS-substrates^52–54^. The difference in *K_m_* argued against the hypothesis that the same active site counts two rounds of reaction before shutting itself off. It further confirmed that the fast- and slow-acting components co-exist in the heterogeneous holoenzymes. Because of the slow OFF-rate (*k_OFF_* < 1/72000 *s^-1^*; at the AGGG position)^52^, *K_m_* ~ *k_OFF_* / *k_ON_* could provide an upper-limit for *k_ON_* of ~ 5.0 x 10^2^ *M^-1^s^-1^*, suggesting that the reaction of the holoenzymes would be slow if the telomere concentration is at the nM level in cells.

### Two active sites in a dimer bind substrates in a sequential manner and act in tandem

Because our two step R3-binding can separate the dimeric enzymes in E1 state, we next ask if the two active sites in a dimer can function at the same time. The kinetic differences between the slow and fast-components could be explained by different Markov models --- a two-step sequential model in a dimer, a parallel model in a dimer or two different monomers (fM_1_ and sM_1_), and a sequential model with a delay in a dimer (**Fig. 4G**). Here the parallel model accounted for either *two* different types of monomeric enzymes or *one* type of dimeric enzymes harboring two distinct active sites. Using MATLAB programs to derive numerical solutions to each model (supplementary information; red-lines in **Fig. 4E-F**), we estimated two kinetic constants from these models, ~0.4 min^-1^ and 0.01 min^-1^ for the fast and the slow components, respectively (**Fig. 4H**). All Markov models fitted the time-lapse data reasonably well. The sequential model with a delay worked slightly better than the others.

To experimentally test the sequential or parallel actions of the two active sites in the dimeric enzymes, we examined whether the two sites interfere with each other during the DNA substrate-binding step, i.e. whether the DNA substrate binding to the first site affects the binding to the second one. The parallel model of one dimer or two different monomers would support simultaneous binding of two substrates to two active sites (**Fig. 4G**). The different monomers must act in parallel. In contrast, a sequential model of a dimer would indicate that the enzyme extends one substrate at a time and only after the first site finishes its catalysis can the second site performs its reaction.

We would like to test whether the dimers can act as in a sequential model. We used the tethered recombinant telomerase holoenzymes, which were physically separated from each other by at least 50 nm on the surfaces of streptavidin-coated beads so that on average, two neighboring separate enzyme complexes were physically independent of each other (**Fig 5A**). Biotinylated telomerase on the streptavidin-coated beads was first incubated with the TS primers (DNA substrates) which would be stabilized at the GGG-position and would stay bound when there was no catalysis ^32^. The saturating concentration of TS primers would fill in all accessible active sites (sM_1_, fM_1_, E_1_ and **E**_0_). After the first round of extension the products were separated from the tethered enzymes within ~20 minutes, which was much shorter than 10 hours such that only a very small fraction (<4%) of the bound substrates in the slow-acting sites (sM_1_, or in either E_1_ or E_0_) that were not catalyzing would dissociate. During this round, monomeric fM_1_ would all become inactive, a good fraction (82%) of monomeric sM_1_ or dimeric E_1_ would remain bound with the products, and all E_0_ dimers would become E_1_. If both active sites of E0 were bound with the substrates, then at the end of the first round, all leftover active sites would be occupied by the DNA-substrates.

**Figure 5:**
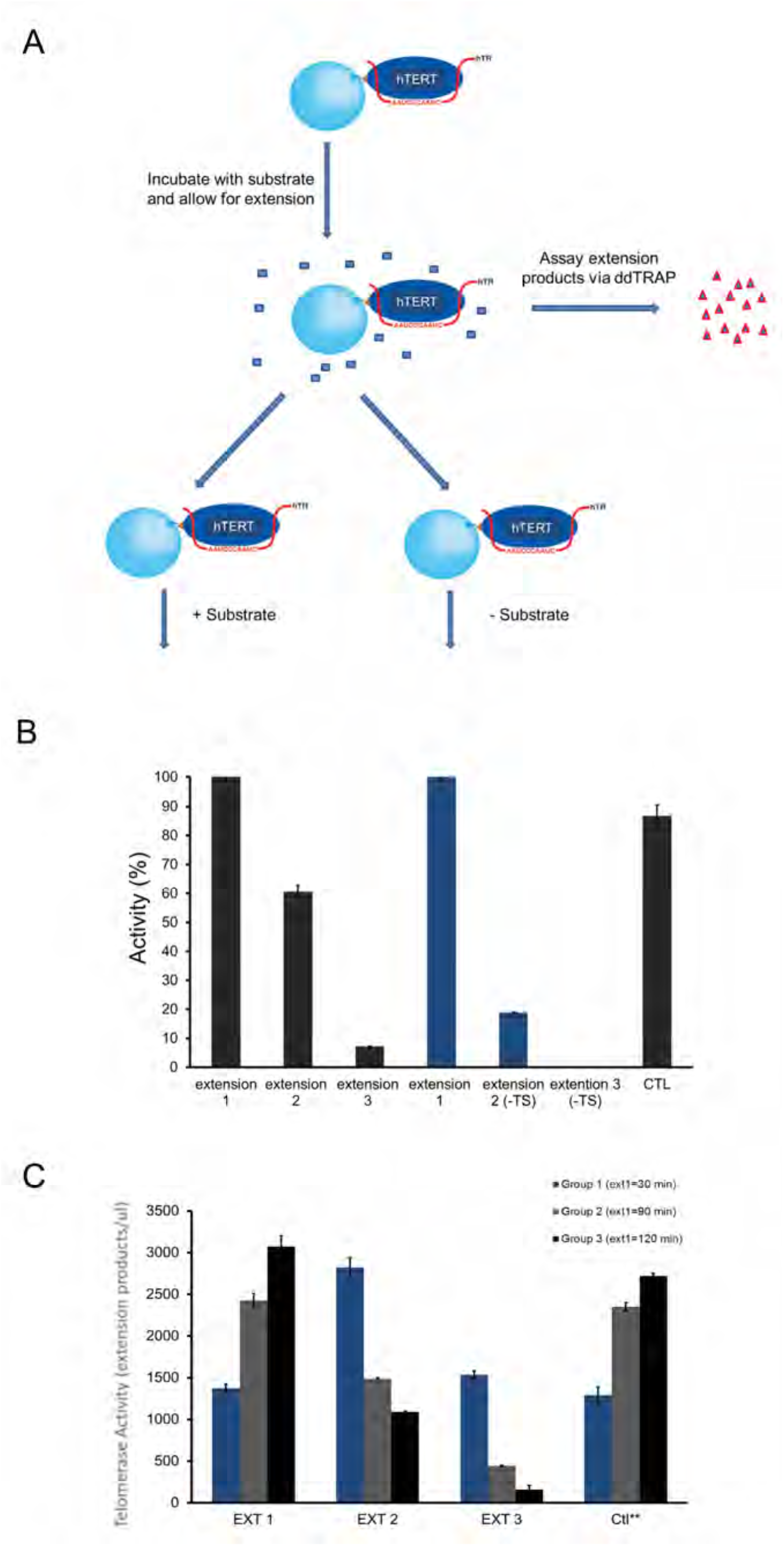
Sequential binding of two active sites to substrates suggests a sequential kinetic model for dimeric enzymes. ***(A)*** Schematic representation of tethered telomerase being tested for whether two active sites are saturated by substrates at the same time. ***(B)*** ddTRAP assay on samples obtained from experiments as designed in panel A. Telomerase extension products were collected and equal amounts were used for the ddTRAP assay. Error bars: *s.d*. (n=3). ***(C)*** ddTRAP assay of tethered telomerase performed at varying times for extension 1 only. Blue, grey, and black bars represent 30, 90, and 120 min respectively. Extensions 2 and 3 were all performed for 2 hours. “Ctl**”samples were time delayed samples that were left at room temperature for five hours before starting the extension reaction for a duration of 30, 90 or 120 min. Error bars: *s.d*. (n=3).

We can test if the above prediction is true by conducting the second round of reactions in two different conditions. For the second round of the reaction, the enzymes on the beads were split into two equal aliquots and performed reactions in an extension mixture with or without fresh TS primers (“ctl + TS” and “–TS” in **Figs 5A & 5B**). If all slow-acting active sites were saturated during the first round of substrate binding, >96% of bound substrates would not have enough time to dissociate during the fast separation of tethered enzymes from free TS primers and products, the second round of extension reaction from the bound-primers should produce approximately the same amount of products, regardless of whether or not fresh primers were added. However, our data showed the opposite. In the second round of reaction the tethered enzymes with no fresh substrates (“-TS”) exhibited a significant drop (~75% less) in extension products, when compared to the control sample (**Fig. 5B**). The significant reduction (44% of the total activity in the first round) in the yield of products during the second round must have come from the *lack of bound substrates* in a majority of the slow-acting active sites of the dimeric enzymes (E_0_) during the first round of reaction, and fresh substrates were necessary for these empty sites to perform their reactions. The residual 16% [Extension 2 (-TS)] could come from the majority (82%) of slow active sites in dimers (E_1_) or slow monomers (sM_1_) that were in the middle of their reactions or from a small fraction of active sites (< 6% based on the extension 3 (+TS) in **Fig. 5B**) that completed their reactions but did not release their products quick enough.

Our observation was not caused by the degradation of the bound TS primers because we used DNAse/RNAse free solutions and nuclease inhibitors in our buffers, and the amount of DNA substrates overwhelmed the active sites of the telomerase by at least nine orders of magnitude. Thus, the only sound explanation was that a significant fraction of slow active sites (75% for Extension 2, equivalent to ~44% of total active sites in Extension 1) were inaccessible to the substrates during the first round of reaction, and thus were *empty*. Based on the data in **Fig. 5B**, we estimated that at the beginning of the first round of reaction, the enzyme mixture contained approximately 44% E_1_ dimers, 36% fM_1_ monomers, and 20% (sM_1_ monomers + E_1_ dimers). In an E_0_ dimer, TS primer-binding to its fast-acting site negatively impacts on the binding of a second primer to the slow active site. This represents a strong negative cooperativity between the two active sites in the E_0_ dimer.

Because of no physical contact between neighboring telomerase complexes on the streptavidin-coated beads, the strong negative cooperativity is only possible when the two types of active sites are within a dimeric telomerase holoenzyme, not from two separate enzymes (**Figs 5A-B, 4G**). Further, negative cooperativity in substrate-binding suggests that a significant fraction (~44%) of E_0_ dimers exist in the enzyme mixture separated from cell lysates even though the new intermediate-resolution cryoEM structure reveals a monomer. The negative-cooperativity also explains the observation that the two active sites exhibit different affinities to the same substrates (**Figs 4C-4D**).

As depicted in **Fig. 4G**, a dimeric enzyme with two different active sites may have three different states. The dimers in cell lysates thus belong to three subgroups --- E_0_, E_1_ and E_2_. In the first round of extension, the pristine enzymes (E_0_) were converted to E_1_ (**Fig. 1A**), and a small fraction of E_1_ changed into E_2_ (**Figs 1A, 4A & 5A**). In the second round, the leftover E_1_ enzymes became inactive E_2_, explaining why the flow-through fraction, the wash ones, or the Elution 2 fraction in **Fig. 1B** had no activity. A sequential model (with or without a delay) is therefore supported by experimental data (**Fig 5B**) and can explain the kinetic property of the dimers (**Fig. 4G**). Between the two monomeric forms, the slow sM_1_ enzymes behaved the same as the E1 and contributed partially to the products seen in Extension 2 (-TS) and the fast fM1 enzymes (36% in the mixture) turned into inactive M_2_ (**Fig. 5B**). The ratio of monomers vs. dimers estimated from our results is thus similar to what was derived by single molecular imaging ^31^.

To further test the sequential model for the fraction of dimeric telomerase, we varied the incubation time for the first extension on the tethered enzymes. For this series of experiments, the same amounts of enzymes were allowed to extend the TS primers for 30, 90, or 120 min in the first extension, respectively. Afterwards, the enzymes were separated from the products and subjected to an equal extension time (90 minutes) for the second and the third round. Based on its time-constant, the fast-acting sites became completely inactive after ~10 minutes (Round 1). The fixed amount of time in the three rounds within the regime of a sequential models predicted that the total products from the three rounds would be similar for three groups. Group 1 (30 min extension in round 1, **Fig. 5C**) had the lowest activity in the first round, but greater activity in both the second and third rounds (**Fig. 5C**) as compared to group 2 (90 min ext 1) and group 3 (120 min ext 1) samples. The enzymes in group 3 with sufficient time during the first two rounds showed almost no activity in the third round because of the catalysis-dependent shutoff. The time-dependent catalysis of the telomerase complex followed the prediction from the models in **Fig. 4G** with substantial amounts of dimer enzymes. Because of the processive reaction in each active site, we name the catalysis-dependent shutoff of human telomerase “*single-run catalysis*”, instead of single-turnover (**Fig. 4G**).

A significant fraction (~44%) of functional enzymes in the cell lysates were dimeric E_0_. Our analysis accounted for the existence of monomeric enzymes, which followed single-exponential kinetics (**Fig. 4G**). Combination of the sequential model for the dimeric holoenzymes with the parallel model for two monomeric forms was thus able to fit all experimental data very well (**Figs 4** and supplementary **Fig. S7**). It is thus not necessary to biochemically purified the telomerase monomers from the dimers for us to deduce the fundamental property of the enzyme. A simple assumption of the fM_1_ and sM_1_ following the kinetics for fast- and slow-acting sites was sufficient for reaching the biophysical understanding (**Fig. 4H**).

### Intracellular telomerase-activating factors (iTAFs) turn on inactive telomerases

Allosterically, the inactive enzymes probably reside in a stable conformation that does not switch back to the active state due to an energy barrier or due to the missing of accessory factors that might dissociate during the middle of the extension reaction. There are three potential destinies for inactive enzymes inside a cell: being recycled into the active form, being degraded or being stored in cells as a reserve. Telomerase is known to be active in dividing cells in every cell cycle. This led us to the question whether it is possible to turn on the inactive telomerase.

We first tested the cell lysates from a telomerase-negative cell line, BJ fibroblasts, and used ddTRAP to quantify the fraction of enzymes that were reactivated (**Fig. 6A**). The tethered enzymes on the streptavidin-coated beads were made fully inactive after three rounds of extension reactions (in E_2_ or M_2_ states in **Fig. 6B**). Afterwards the beads with inactive enzymes were mixed with the BJ cell lysates and the extension reaction mixture for 2 hours before the extended products were counted using ddTRAP. To our surprise, the cell lysates from the telomerase-negative BJ cells were able to reactivate a significant fraction of inactive enzymes (as exemplified in **Fig. 6B**). As a negative control, the BJ cell lysates had no activity (**Fig. 6B**). These results clearly demonstrated that the BJ cells contain intracellular telomerase-activating factors (iTAFs) that can reactivate the inactive holoenzymes (E_2_ and M_2_). The same iTAF activity was detected in other types of telomerase-negative proliferating cells, such as SAOS-2 and SKLU-1 (**Fig. 6F**).

**Figure 6:**
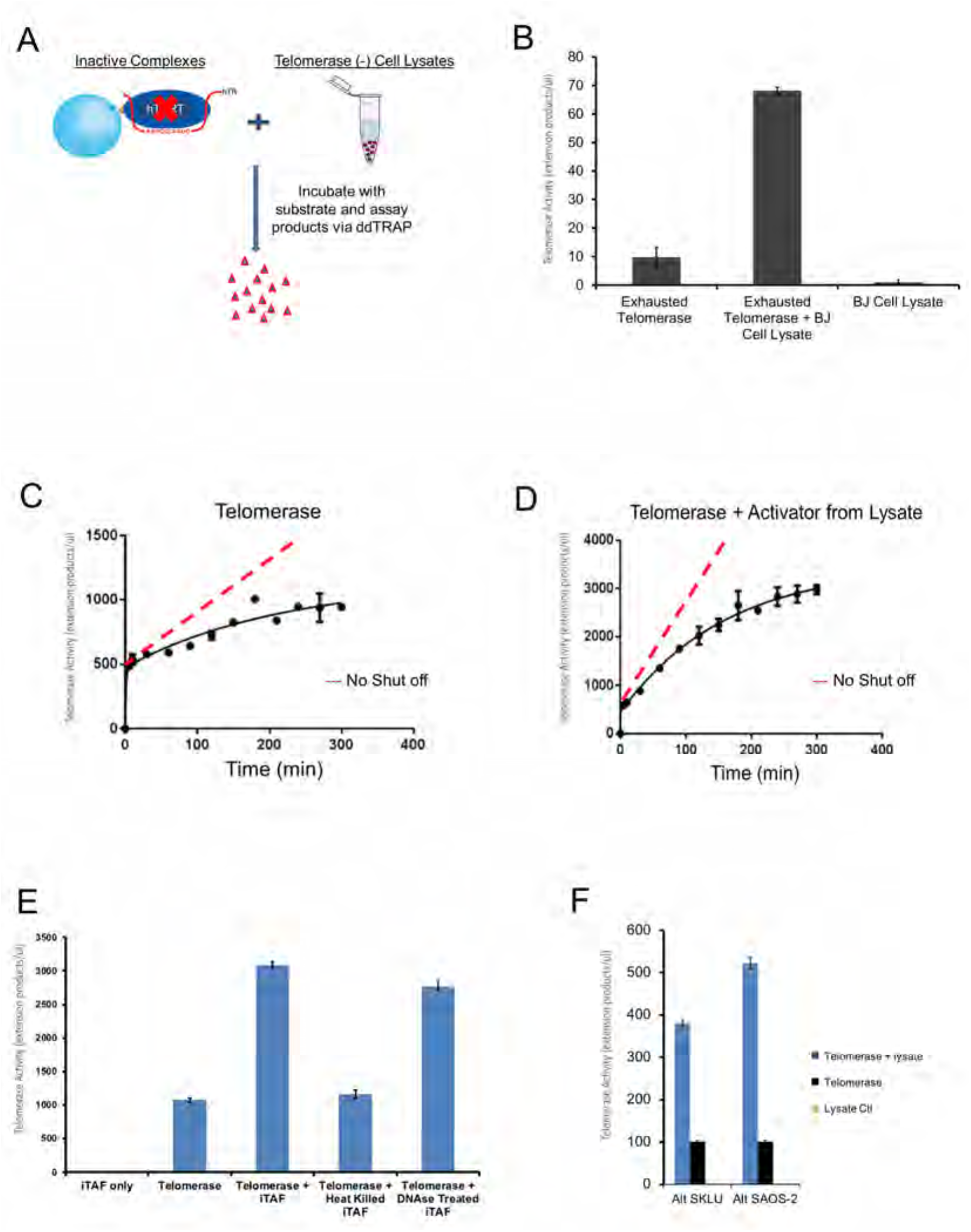
iTAFs reactivate inactive telomerase holoenzymes. ***(A)*** Schematic representation of catalytically exhausted tethered telomerase enzymes (E_2_ or M_2_) mixed with telomerase-negative cell lysate (of BJ cells) to test enzyme reactivation. ***(B)*** Telomerase reactivation assay performed using ddTRAP. Error bars: *s.d*. (n=3). Inactive telomerase treated with the BJ cell lysate showed a 7-fold increase in activity compared to the background readout from control enzymes. BJ cell lysate shows no telomerase activity. ***(C)*** & ***(D)*** Time courses for the tethered telomerase before and after treatment with the iTAFs (from BJ cells). The continued lines were fitted with the sequential model with a short delay. After reactivation, the slow-acting enzyme is dominating (75%). Error bars: s.d. (n=3). The dashed lines represent the amount of products if the active sites remained continuously active. ***(E)*** iTAF fractions were heat-inactivated or treated with the DNAse before being incubated with the inactive tethered telomerase. The background activity was from the tether E2 enzymes with residual activity. Other samples were normalized against the control. Error bars: *s.d*. (n=3). ***(F)*** Cell lysates of SAOS-2 and SKLU-1 contain iTAF activity. These are two alternative lengthening telomere (ALT) cells lines that have no telomerase activity.

When partially purified telomerase holoenzymes were incubated with iTAFs, the time-lapse experiments showed that the activation by iTAFs increased the amount of the fast-acting sites (E_0_ or fM_1_) by only a small fraction (~25%), but boosted the slow-acting active sites (E_1_ or sM_1_) much more significantly (~480%) (**Fig. 6C vs. 6D**). The control telomerase (**Fig. 6C**) had roughly equal amount of fast and slow sites. The (re)activated enzymes were dominated by slow-acting sites (E_1_ or sM_1_), suggesting that the iTAFs preferentially switched the inactive enzymes into the E_1_ or sM_1_ states. Furthermore, the fast active sites in the reactivated enzymes displayed a slightly faster kinetics, which remains comparable to that of the native enzymes. These data further demonstrate that the two kinetic components reflect the intrinsic properties of the native, functional telomerase holoenzymes (supplementary **Fig. S7**).

The iTAFs are proteineous components. We were able to separate iTAFs from the telomerase fractions in a glycerol gradient. The iTAFs heated in a boiling temperature lost their activity in reactivating the inactive enzymes (**Fig. 6E**). DNAse treatment of iTAFs had almost no effect on their activity (**Fig. 6E**). When the cell lysates were fractionated and eluted in a Superdex 200 column, the iTAFs were found to be equivalent to an ~150 kDa globular protein (supplementary **Fig. S8**). Further experiments are needed to identify the iTAFs and characterize their activity on the holoenzymes.

## DISCUSSIONS

### A kinetic model for catalysis-dependent inactivation and iTAF-mediated reactivation

Our results show that in the telomerase mixture of both dimers and monomers inside a human cell, a dimeric telomerase containing both fast- and slow-acting active sites and there are two isoforms of monomeric enzymes harboring similar fast and slow active sites, respectively (**Figs 4A-D**). The two types of active sites exhibit different affinities for the DNA substrates. The negative cooperativity between them happens within a dimeric enzyme and makes the slow site inaccessible to substrates when the fast site is in catalysis. A dimeric E_0_ enzyme may undergo a symmetry breakdown when one of its active sites is bound with a DNA substrate, and its two sites act in tandem. The monomers do not have negative-interactions between them (**Fig. 5B**). More importantly, both types of active sites perform single-run catalysis (**Fig. 7A**), and iTAFs can turn the inactive enzymes (E_2_ or M_2_) into different active states (**Fig. 7B**). Our results can be incorporated into a kinetic model in **Fig. 7A**, where one active site goes through one round of processive extension reaction and becomes inactive after it falls off the product (**Figs 1A-B**). The inactive site can be turned back on (step IV; recycle). The switching of the enzymes between active and inactive states is further diagramed in **Fig. 7B**. Both the endogenous telomerase holoenzymes and the recombinant ones exhibited similar kinetic property under different conditions. We therefore propose that the use-dependent loss of activity and the iTAF-dependent gain of activity represent fundamental properties of human telomerase and provide an intrinsic ON-OFF control of its activity in human cells.

**Figure 7:**
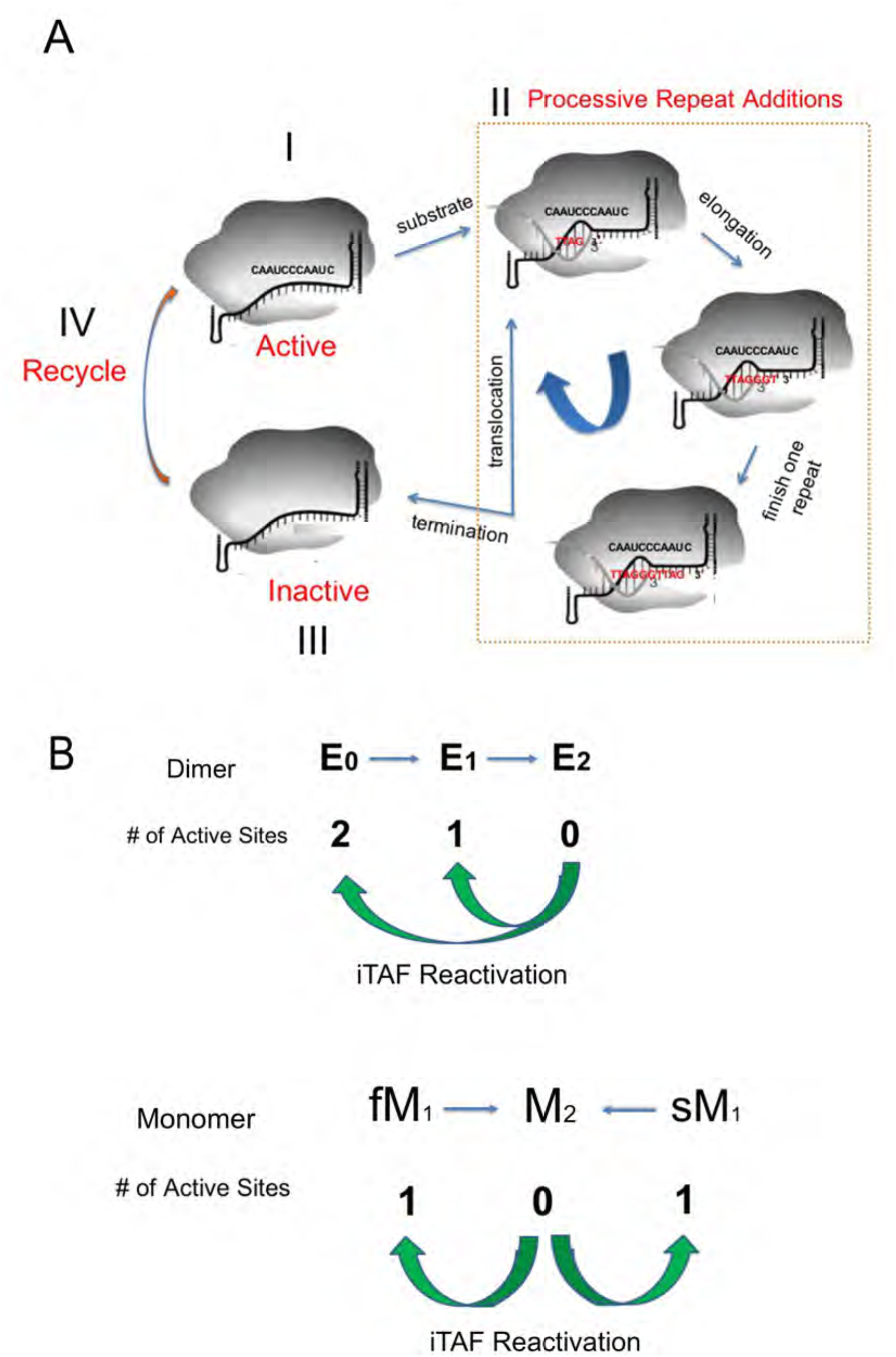
Kinetic ON-OFF control of human telomerase holoenzyme. ***(A)*** Each active site undergoes catalysis-dependent shutoff. During the processive catalytic reaction, after each translocation of the newly added repeats the enzyme has a frequency in falling off the substrate, which results in the shutdown of the active site. The iTAFs can reactivate the active site. The dashed line box denotes the processive catalysis of the telomerase activity. ***(B)*** A sequential model for the two active sites of the human telomerase holoenzyme and the parallel action of two different monomeric enzymes. Newly assembly dimeric enzymes (E_0_) have two active sites. E_1_ has its fast-acting site shutoff, and E_2_ has no activity. M_1_ has one active site and M_2_ none. The iTAFs can switch the inactive sites into active states through E_2_ → E_1_ or E_2_ → E_0_ with E_2_ → E_1_ dominating the reactivation or through M_2_ → M_1_. The ratio between the switches of M_2_ to fM_1_ and to sM_1_ is unclear.

The sequential action of the two sites in a dimeric telomerase (**Fig. 7B**) suggests that a dimeric enzyme does not act on two chromosomal ends (say homologous chromosomes) at the same time ^43, 48, 55^. Two monomers can act on two telomeres simultaneously. The total time for complete exhaustion of telomerase activity would be ~300 minutes, shorter than the average duration of the S-phase of human cell cycles (6-8 hours). For quick dividing cells, the S-phase might be < 30 minutes such that only the fast-active sites (E_0_ and fM_1_) are suitable for fast chromosomal replication. Under such conditions, extra copies of mature telomerase holoenzymes would be needed to avoid critically short telomeres. A combination of fast and slow active sites in dimers and monomers would be sufficient to match the time needed for telomerase to finish its catalysis with the duration of the S-phase so that the telomerase activity can be maximally utilized. This might be one of the reasons why the telomerase holoenzymes have two types of active sites with contrastingly different kinetics. This scenario may still allow the reactivation of fast-acting sites to perform extra rounds of reactions when cells need them to ensure timely completion of the S-phase. This mechanism indicates a possible role of the iTAFs in regulating the S/G2 transition.

### Single-run catalysis of human telomerase

When counting the extension products by ddTRAP with single digit accuracy, we could avoid complications from the average length of extension products, the differences in catalysis rates, and the possible delay between the moment the fast site in a dimer is turned off and the time point when the slow site becomes accessible to a new substrate (**Fig. 4G**). Using the substrate pull-down to select only active enzymes in the single-turnover setting (**Figs 1A-1B**), we avoided the often observed mismatching between a larger amount of enzymes (or hTERT) and a smaller number of extension products in the enzyme assays ^34^, which indicates a significant fraction of inactive enzymes. The similar distribution of the products of different lengths in the Input 1 and that in the Elution 1 (E_1_ or slow M_1_) in **Fig. 1B** from the endogenous telomerase suggests that the processive additions of repeats to substrates are similar between the slow and the fast active sites (**Fig. 7A**). The main differences between the fast and slow-acting sites are thus due to other factors, such as accessibility to substrates, initiation of the successful extension reaction, pause duration after the complete extension of each repeat, etc. Singe molecule enzymology would be suitable for determining the detailed differences between these two active sites ^44, 56, 57^, even though the slow sites might be difficult to study and the dimeric enzymes might fall apart *in vitro* and the enzymes remain heterogeneous in stoichiometry and conformation.

The single-run catalysis may not be limited only to human telomerase ^58, 59^ because the monomeric enzymes seen in other organisms may have two different isoforms. It is tempting to speculate a similar mechanism in yeast and Tetrahymena even though they vary in TERT and TR size and in complex stoichiometry. The single-run catalysis also occurs to other ribonucleoprotein (RNP) complex, for example, CRISPR/cas9 ^60^, suggesting a more universal mechanism among RNP enzymes. A generic mechanism of catalysis-dependent turn-off and iTAF-dependent turn-on of the telomerase holoenzymes in other species will likely reveal a more general role of the single-run catalysis in achieving tight regulation of the telomerase activity.

### The iTAFs differ from other proteineous factors that regulate telomerase activity

The iTAFs are proteineous factors that can be fractionated. They are the first group of factors that directly control the recycling of human telomerase, and are mechanistically different from the recruitment factors in the shelterin complex, the hTR binding factors as well as the factors involved in the assembly of nascent active enzymes ^61^. It is still early to say whether they function at the same location and / or time as the recruitment and/or activation of the telomerase at the shelterin complex ^61, 62^. Nor are we able to explain why iTAFs are found in telomerase-free cells. Future study will be needed to identify the iTAFs and verify their functions in both telomerase-positive and -negative cells.

With the slow kinetics and the stability of the telomerase holoenzymes within 5-8 hours at room temperature, we have not been able to determine how many times each telomerase can be turned off and turned back on before its breakdown. The high stability of a holoenzyme in the nuclear environment may allow multiple OFF-ON cycles before it becomes defective and is marked for degradation. More broadly, whether the iTAFs function in controlling cell cycle and what other functions they might have in cell proliferation and cell ageing are interesting questions for the future.

### Single-run catalysis as a built-in brake for human telomerase

Similar to single-turnover enzymes, catalysis-dependent inactivation of human telomerase offers an exquisite regulatory mechanism. There are at least two layers of control. The first is that the fast-acting sites are quickly exhausted, ~40 times faster than the slow sites. The second is that the iTAFs may be regulated through different pathways.

The single-run catalysis explains the need for telomere extension to happen in focused areas in the nucleus. With each active site acting once on one telomere in a co-replicational manner in the S-phase, ~92 active sites are needed, that is equivalent to 46 copies of dimeric telomerase holoenzymes in E_0_ state to satisfy the need of extending all telomeres. If these enzymes are randomly distributed in a nucleus of ~5 microns in diameter, the average concentration of the active sites is ~ 0.1 - 0.2 nM, two orders of magnitude lower than the measured *Km* for these sites (**Figs 4C-D**). To solve this mass-action problem, the telomerase molecules may be concentrated in a small area (< 0.5 microns) such that their local concentration would be ~200 nM, sufficient to support productive collisions between telomerase active sites and individual telomeres. Similarly, the telomeres of different chromosomes may be looped from individual chromosomal territories into the telomerase-concentrated regions, the telomere processing centers, so that the local concentration of the substrates (uncapped telomeres) also can approach ~200 nM, which would be sufficient for ensuring a sizeable volume of extension reactions within a short period of time.

In conclusion, the native human telomerase has two kinetically distinct types of catalytic sites that manifest varying affinities for the telomeric substrates. The two sites in dimeric enzymes show negative cooperativity and act in tandem by sequentially becoming accessible to the DNA substrates. The active sites in monomeric enzymes function independently. Both types of active sites function as a single-run enzyme because they undergo catalysis-dependent shutoff. Inactive enzymes can be reactivated by iTAFs from certain cells. Identification of the iTAFs and their roles in regulating telomerase activity await future studies.

## Acknowledgements

The authors are indebted to members of the Jiang laboratory and those of the Shay/Wright laboratory. The results in the manuscript represent part of the PhD dissertation work of M.E.S., which was submitted to the UT Southwestern and University of Texas at Arlington in August of year 2016. Three members of the thesis committee, Drs. Hamid Mirazei, Robert Eberhart and Michael Cho, provided insightful advice and comments. Drs Sixue Chen and Jin Koh in the Proteomics Core at the Interdisciplinary Center of Biotechnology Research (ICBR) of UF offered technical support. This work was mainly supported by CPRIT (RP120474 to Q.-X.J.), and partially by NIH (R01GM111367 & R01GM093271 to Q.-X.J.), CF Foundation (JIANG15G0 to Q-X.J), Welch Foundation (I-1684 to Q.-X.J.) and the startup funds (to Q.-X.J) from UF. Some experiments reported here were performed in a laboratory constructed with support from NIH (grant # C06RR30414 with Dr. Jerry Shay as the PI). A.T.L. was funded by a K99/R00 Pathway to Independence award from NCI (grant number KCA197672A).

## MATERIALS AND METHODS

Cells used for preparing active telomerases were either H1299 lung adenocarcinoma cells or super H1299 overexpressing both hTR and N-terminally biotinylated hTERT. BJ human fibroblasts and other cells were all prepared using proper procedures. Details are in the supplementary information.

RT-qPCR (reverse-transcribed quantitative polymerase chain reaction) of hTR was performed by RNA purification, reverse transcription to produce cDNA, and qPCR using proper primers against hTR cDNAs.

Gel-based TRAP assay was performed as described before ^63^. Digital droplet PCR (ddPCR)-based TRAP assay (ddTRAP) quantified the individual extension products by running ddPCR reactions in individual droplets containing either one or none product, and detecting the positive ones based on fluorescence. Detailed procedure can be found in Ludlow et al. ^47^. ddTRAP counts the individual extension products in the range of 0-15,000 with high accuracy and reproducibility.

SDS-PAGE analysis and western blot detection of protein components were performed based on published procedures ^64^.

Partial purification of recombinant telomerase was done in four steps. 200 −500 million super H1299 cells were pooled together, and lysed in a 1.5% CHAPS buffer. The lysate was cleared by centrifugation and then fractionated in a continuous glycerol gradient. The active fractions were pooled together and incubated with monomeric avidin beads (Pierce). After the beads were washed with a buffer, the active enzymes were eluted with 2.0 mM D-biotin. The active fractions were pooled together and incubated with SPFF beads (GE Health Science) and eluted with 0.2 - 1.0 M NaCl. The eluted enzymes were further fractionated by size-exclusion chromatography in a Superose 6 column (SEC).

5’-biotinylated telomeric repeats [(TTAGGG)_3_; R3] were loaded to streptavidin-coated Dyna-beads. After complete wash to remove free R3, the R3-conjugated beads was used for single-step pull-down of human telomerase holoenzymes as described before ^32^. dATP/dTTP were used to elute the bound enzymes from the beads. Apyrase treatment was introduced to remove the nucleotides when two such pull-down steps were performed in tandem.

Streptavidin-coated magnetic Dyna-beads (MyOne T1, Invitrogen) were used to present the biotinylated telomerase holoenzymes for multiple rounds of extension reactions. Based on the surface area of each bead and the amount of streptavidin molecules used for cross-linking reaction, the average distance between neighboring streptavidin molecules would be at least 50 nm. A magnet was used to achieve quick separation of the tethered enzymes and the reaction products in less than 3 minutes. Similarly, protein A/G-coated magnetic beads were used to present anti-myc antibodies, which can recognize the myc-tags introduced to the N-terminus of the recombinant hTERT.

Partial purification of intracellular telomerase-activating factors (iTAFs) from cell lysates followed a four-step procedure. After cell lysis, the cleared lysates were fractionated in a continuous glycerol gradient. The fractions containing iTAF activity were pooled and concentrated before being run in a Superdex 200 gel-filtration column. The active fractions from size-exclusion were loaded into a Mono-S column or SPFF for further purification. The active fractions were tested and subjected to further kinetic analysis.

The on-line supplementary information contains more details for each step.

